# Birdsong modification with food reward

**DOI:** 10.64898/2026.05.27.728162

**Authors:** Franziska Heubach, Lena Veit

## Abstract

Songbirds, such as Bengalese finches (*Lonchura striata domestica*) produce syntactically organized vocal sequences, composed of individual syllables strung together in variable order. Adult birds can learn to modify transition probabilities between syllables through sequence modification training: when punishing one transition with manipulated auditory feedback, birds will gradually reduce the targeted transition. This protocol is thought to rely on a circuit for vocal-auditory monitoring of the bird’s own song. Despite the overwhelming usefulness of food rewards for other kinds of trained animal behavior, reinforcing birdsong features with food rewards has remained elusive, possibly because the slow timescale of food reward does not match the fast and precise vocal-auditory feedback loops underlying learned birdsong. Here, we use second-order conditioning to selectively increase the frequency of target transitions in song with food rewards. Bengalese finches learned to associate a delayed primary reinforcer (food delivery by an automated feeder after song ends) with a secondary reinforcer (a short click sound). We then reinforced specific target syllables with the click during ongoing song, leading birds to selectively increase the frequency of the targeted syllable transitions. Learned changes were specific to the target and evident in catch trials without reinforcement, indicative of a learning process. Our results demonstrate that song control circuits can learn from different feedback modalities, including learned associations with food reward.

## Introduction

Reinforcement learning is a fundamental process that enables both humans and animals to adapt diverse behaviors based on experiences. The consequence of a behavior can act as a reinforcer, and a positive outcome can increase the probability of showing the behavior in the future [1–4].

Birdsong is a rare example of vocal learning in animals, with well-established parallels to hu-man speech learning [5,6]. Song learning by juvenile birds relies on precise auditory feedback-dependent learning processes that lead to increasing similarity between juvenile song syllables and tutor models [7–10]. It can be understood within the framework of reinforcement learning [11–13], with auditory similarity to the tutor song, rather than external reward, acting as an internal performance measure [10,14]. Even after the initial period of song acquisition, adult birds continuously rely on auditory feedback and learning processes to maintain their song [15–17].

Despite the overwhelming success of primary reinforcers like food or water reward to modify and shape other types of animal behavior [4,18], song modification with positive reinforcement has remained elusive. Adult birds can be trained to modify specific aspects of their song such as syllable structure, timing, or sequencing to avoid negative consequences in the form of manipulated auditory feedback [16,17,19–21]. In these cases, a precisely timed auditory mismatch is created between an aversive white noise (WN) stimulus and the bird’s expectation of its own song, which acts as a basis for reinforcement learning [13,22,23]. Is song modification learning possible exclusively through auditory mismatch? The highly specialized song production circuit might be uniquely adapted to processing perceived auditory error [12,19,22,24–26], and birdsong as a courtship behavior may underlie ‘biological constraints’ on learning [27,28], limiting its associativity with rewards from other behavioral domains, such as feeding. On the other hand, it may be possible for general reward and error-related cues to guide adaptive changes to song [29].

We hypothesized that targeted song modifications with positive reinforcers might be achieved if temporal precision of feedback matches the timescale of song. We trained adult male Bengalese finches (*Lonchura striata domestica*) to adaptively increase rare aspects of their song using positive reinforcement with food rewards. We developed a new setup that achieves the temporal precision required for song modification learning by using a short auditory stimulus as a secondary reinforcer during ongoing song [3,18,30], and delivering food after song ends. Bengalese finch song consists of acoustically discrete syllables that occur in variable sequences [31–33]. Transition probabilities between syllables have previously been reduced with aversive auditory feedback [17,34–36] but it has not been possible to increase rare transitions in a targeted way. We show that rewarding a specific syllable after a branch point, where one syllable can be followed by two or more distinct syllables [32], dramatically increased its transition probability. Rewarding specific numbers in repeat phrases, where the same syllable type is re-peated [31,33,37,38], shifted repeat distributions toward the rewarded repeat numbers. Our results show that sequence modification learning in songbirds is possible using positive reinforcement with food reward, and precise temporal control over the specific transitions can be achieved through second-order conditioning.

## Results

### Transition probabilities can be modified using positive reinforcement

We trained four birds to first associate a short click sound as a secondary reinforcer with delayed feeder access in a conditioning phase (Fig. 1A). All birds successfully learned to approach the feeder after the click. We then used Moove [36] for real-time detection of target syllables, and connected its output to an Arduino Uno controlling the click and feeder (see Methods). In the pilot bird (bird 1), we targeted a branch point syllable to test our protocol in comparison to a well-established song modification protocol with aversive auditory WN feedback. We chose the rare branch ‘d’, preceded by ‘bk’, for targeting with the click (Fig. 1B). After several days of training with song-contingent positive reinforcement, the transition probability from ‘bk’ to ‘d’ significantly increased by 339% compared to baseline recordings (Fig. 1C, D; transition probability ‘bk-d’ baseline 5.3%, training 23.1%; Fisher’s exact test: odds ratio = 0.2, p < 0.01). In line with previous studies [17,34,35], transition probabilities returned towards baseline after training (Fig. 1D).

**Fig. 1:**
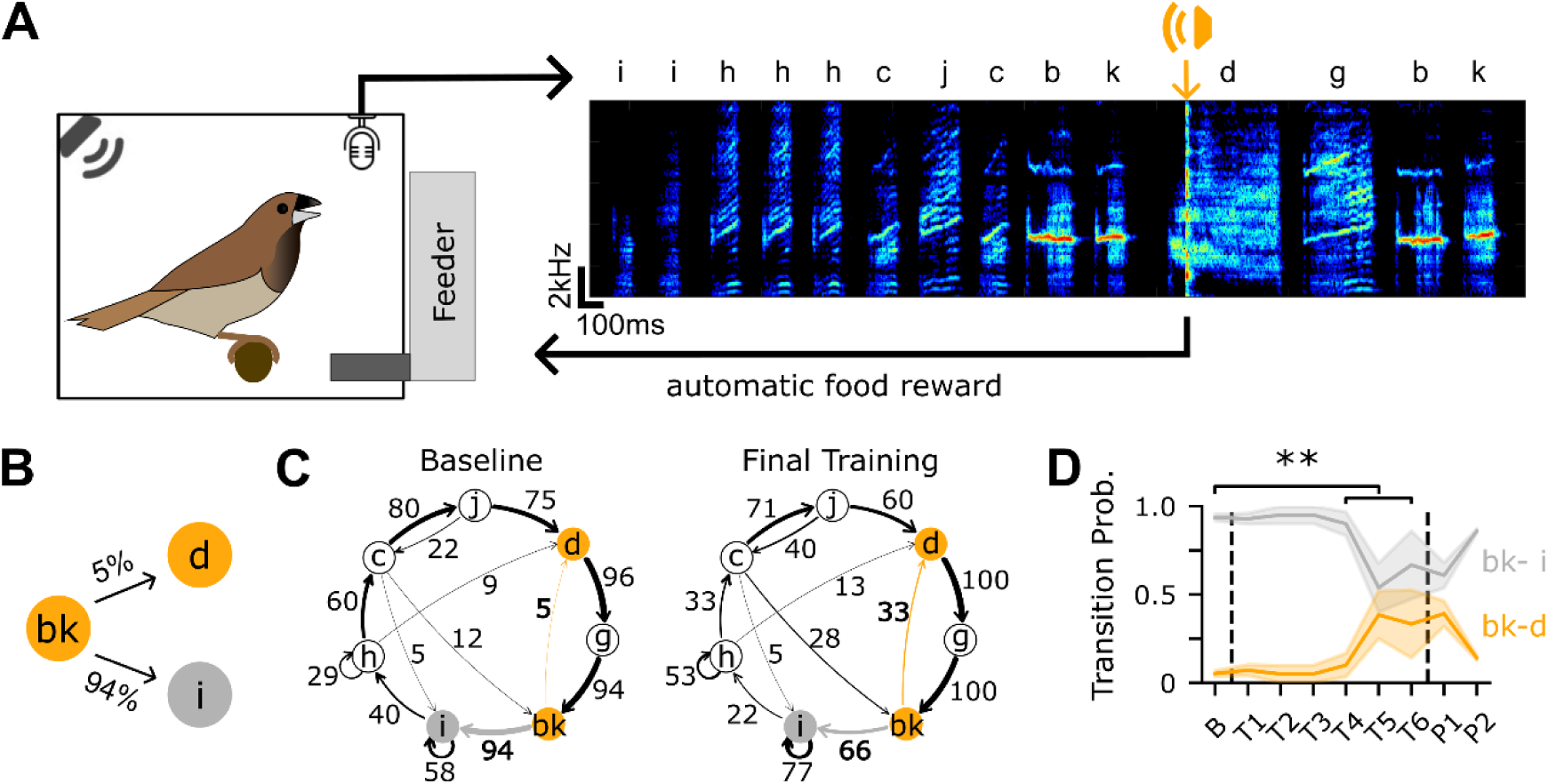
Rare syllable transitions can be increased by positive reinforcement with food re-wards. A) Schematic of the training setup. Example spectrogram of bird 1, with individual syllables labelled by letters. Song is recorded and a click is played following detection of the target syllable onset (orange arrowhead). Food reward is delivered by activating the automated feeder after the song bout ends. B) Branch point targeted in bird 1. Numbers on arrows represent transition probabilities between syllables, the target transition ‘bk-d’ is marked in orange. C) Transition diagram of song during baseline (left) and the final day of training (right). Nodes represent syllables and edges show possible transitions between syllables. Line thickness and numbers on edges represent transition probabilities in percent. The target transition at the branch point is marked in orange, and transition probabilities for the target branch and alternative branch are depicted in bold. Probabilities < 5% are not shown. D) Transition probabilities at the target branch point during baseline (B), six days of training (T1 - T6) and post-baseline (P) recordings; P1 immediately following training, P2 screening on a separate day after approx. one week (see Methods). Dashed lines show boundaries between baseline, training and post-baseline. ** = p < 0.01, Fisher’s exact test.

### Repeat phrases can be extended with positive reinforcement

After establishing that birds can learn to modify transition probabilities at branch points using positive reinforcement, we next investigated whether they could learn to selectively increase syllable repetition in repeat phrases. Repeat phrases are structures in Bengalese finch song where the same syllable type is repeated a variable number of times [31,33,37,38]. We call the number of syllables in one rendition of the phrase its repeat number. We targeted rare repeat numbers at the high end of the naturally occurring distributions (see Methods) with the secondary reinforcer in seven different repeat phrases of three birds (birds 2 - 4). Figure 2A, B shows the spectrogram and repeat distribution for repeat phrase ‘J’ of bird 2. We targeted this phrase at repeat positions 6 and above (Fig. 2B; blue shaded region). After training, this bird significantly increased the mean repeat number of phrase ‘J’ (mean baseline 3.6 ± 0.0, mean training 4.2 ± 0.1; Mann-Whitney U test: U = 35603.5, p < 0.001; all numbers are mean ± s.e.m.) and decreased it again after training (Fig. 2C). Accordingly, the repeat number distribution shifted towards higher repeat numbers during training (Fig. 2B). Across all experiments, the mean re-peat number significantly increased by 24.3% ± 4.5% during training compared to baseline (Fig. 2D; mean baseline 2.7 ± 0.3, mean training 3.4 ± 0.3; Wilcoxon signed-rank test: n = 7, z = 0.0, p < 0.05).

**Fig. 2:**
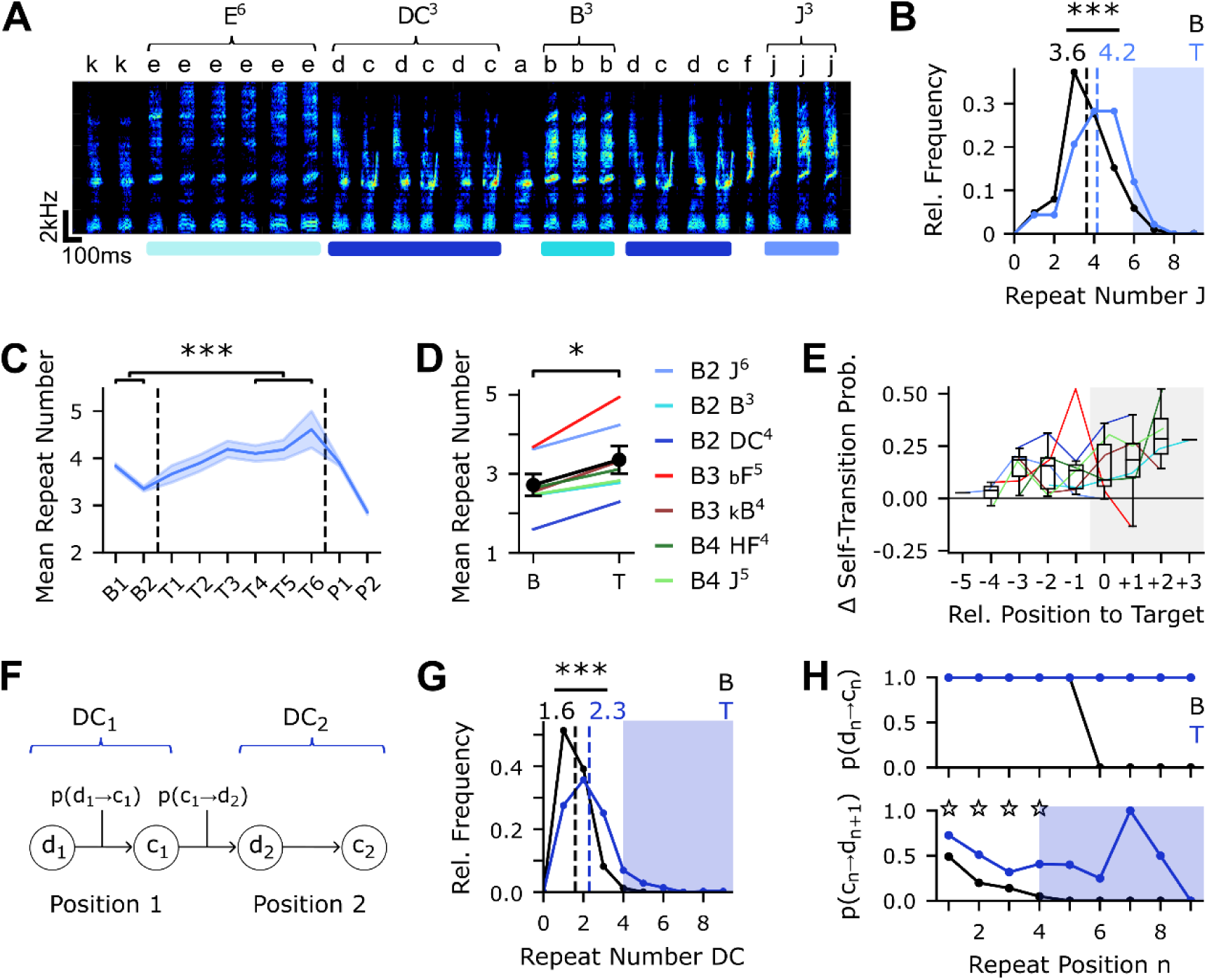
Positive reinforcement can increase repeat number of single-syllable repeat phrases and meta-repeats. A) Spectrogram of an example song bout from bird 2. Repeat phrases are marked by colored bars, and are depicted by a capital letter, with repeat number given in the exponent. B) Repeat distribution of repeat phrase ‘J’ during baseline (B, black) and training (T, blue). Dashed lines show mean repeat number. Blue shaded region depicts rewarded repeat numbers ≥ 6. *** = p < 0.001, Mann-Whitney U test. C) Mean ± s.e.m. repeat number during baseline (B), training (T) and post-baseline (P) for the same repeat phrase shows dynamics of the learning process. Dashed lines show boundaries between baseline, training and post-baseline. *** = p < 0.001, Mann-Whitney U test. D) Mean repeat number during baseline (B) and training (T) for all experiments targeting repeat phrases (n = 7 experiments from n = 3 birds). Birds are indicated by names (B2 - B4), and target phrases for each training are indicated by the phrase and target number, for example J^6^ indicates reward on repeat number 6 and above for phrase J. Black line shows mean ± s.e.m. across birds; * = p < 0.05, Wilcoxon signed-rank test. E) Self-transition probability of the repeat syllable at different positions relative to the target (0). Individual lines depict different experiments, colors as in D) (n = 7). Grey shaded region depicts rewarded repeat numbers (≥ target). F) Example meta-repeat: The chunk ‘dc’ is repeated a variable number of times; repeat number counts repetitions of the entire chunk, e.g. DC_1_. Syllable ‘d’ transitions to ‘c’ with p(d-c) at repeat position n, syllable ‘c’ transitions to ‘d’ with p(c-d), leading to repetition of the chunk from repeat position n to n+1. G) Repeat number distribution of meta-repeat ‘DC’ during baseline (B, black) and training (T, blue). Dashed lines show mean repeat number. Blue shaded region depicts rewarded repeat numbers (≥ 4). *** = p < 0.001, Mann-Whitney U test. H) Transition probabilities in the meta-repeat. Top: within-chunk transition probability p(d-c) for each repeat position during baseline (B, black) and training (T, blue). Bottom: between-chunk self-transition probability p(c-d) for each repeat position. ⋆ = p < 0.001, χ^2^-contingency tests (repeat positions 1 - 3) and Fisher’s exact test (repeat position 4). Higher positions could not be tested because of a lack of baseline data at these repeat numbers.

The shift towards higher repeat numbers could be achieved by uniformly increasing the self-transition probability of repeating syllables. Alternatively, the birds might selectively increase only the relevant transitions leading up to the targeted number. We found that self-transition probabilities tended to increase for all shown positions preceding and following the target (Fig. 2E; mean increase of +16%).

### Higher-order repeating structures can also be extended with positive reinforcement

Songs from birds 2 and 4 contained repeat phrases composed of more than one syllable, such as ‘dc-dc-dc’ in Figure 2A. We called these structures meta-repeats. Meta-repeats were marked by a chunk of multiple syllables (e.g. ‘dc’, see Methods; Fig. 2F), which was repeated a variable number of times. Repeat distributions of meta-repeats resemble the peaked distributions observed for repeat phrases of single syllables (Fig. 2G). We tested whether meta-repeats could be similarly extended with positive reinforcement in a subset of two birds. In bird 2, we re-warded the fourth chunk ‘dc’ and any following chunks. The repeat distribution shifted towards higher repeat numbers as expected (Fig. 2G; meta-repeat ‘DC’: mean baseline 1.6 ± 0.0, mean training 2.3 ± 0.0; Mann-Whitney U test: U = 615110.5, p < 0.001). Across the two experiments, the relative increase of mean repeat number was 31.5% (Fig. 2D; second meta-repeat ‘HF’: mean baseline 2.6 ± 0.0, mean training 3.1 ± 0.1; Mann-Whitney U test: U = 97496.5, p < 0.001). The secondary reinforcer (i.e. the click) was presented on the second syllable ‘c’ of the chunk ‘dc’. As expected from prior results with aversive auditory feedback [17], the fixed transition within the repeating chunk was not modifiable by training (Fig. 2H, top). As with single-syllable repeat phrases, self-transitions of the meta-repeat (i.e. from ‘c’ to ‘d’) increased in all positions preceding the target (Fig. 2H bottom; χ^2^-tests: all p < 0.001, mean increase 110%), and at the target position (Fig. 2H bottom; increase +729.7%; Fisher’s exact test: ratio = 0.1, p < 0.001). Meta-repeat self-transitions following the target also tended to increase, but their absence from baseline recordings did not allow for statistical testing. Thus, the magnitude of changes observed for meta-repeats is similar to that of single-syllable repeat phrases, and changes likewise occur at all tested repeat positions.

### Context-specific repeat phrase modifications

Transition probabilities in Bengalese finch song, including repeat number distributions, can depend on the preceding sequence context [33,39,40]. This suggests that repeat phrases in different sequence contexts may have distinct premotor representations [38,41]. One of the targeted repeat phrases occurred in multiple sequence contexts: In bird 3, we targeted repeat phrase ‘B’ at positions four and above only in the sequence context ‘kB’, but not when it occurred in its other contexts ‘fB’ and ‘iB’ (Fig. 3A, B). As expected, the mean repeat number of phrase ‘B’ increased by 32% compared to baseline in the context ‘kB’ (Fig. 3B, right; mean baseline: 2.5 ± 0.1, mean training 3.3 ± 0.2; Mann-Whitney U test: U = 3354.0, p < 0.01). In contrast, the mean repeat number of ‘fB’ increased only by 15% (Fig. 3B, middle; mean base-line 2.6 ± 0.0, mean training 3.0 ± 0.2; Mann-Whitney U: U = 6243.5, p < 0.01), while the repeat number of ‘iB’ did not increase (Fig. 3B, left; mean baseline 3.1 ± 0.1, mean training 3.1 ± 0.1, Mann-Whitney U test: p > 0.05). Therefore, this bird modified repeat distributions in a context-specific way and exhibited limited generalization to some but not all non-target contexts. Additionally, this bird modified variable transitions in the song to significantly increase the relative occurrence of the targeted repeat context and correspondingly reduced the non-target contexts (Fig. 3C; χ^2^-tests: all p < 0.001). Correspondingly, from the perspective of sequencing decisions at branch points, the probability for ‘k’ to transition to ‘B’ increased significantly during training (mean 16.2% to 37.5%, χ^2^(1) = 43.9, p < 0.001), while the alternate probability ‘k-c’ decreased accordingly (mean 83.8% to 62.5%; Fig. 3D). When considering all transitions in the song, even bigger increases were seen at the transition ‘m-c’, two positions before the ‘k-B’ transition (Fig. 3E). Thus, song sequencing was modified at earlier stages to increase the probability of transitioning to the rewarded context, but the increase in selftransition probabilities was not completely specific to the rewarded context.

**Fig. 3:**
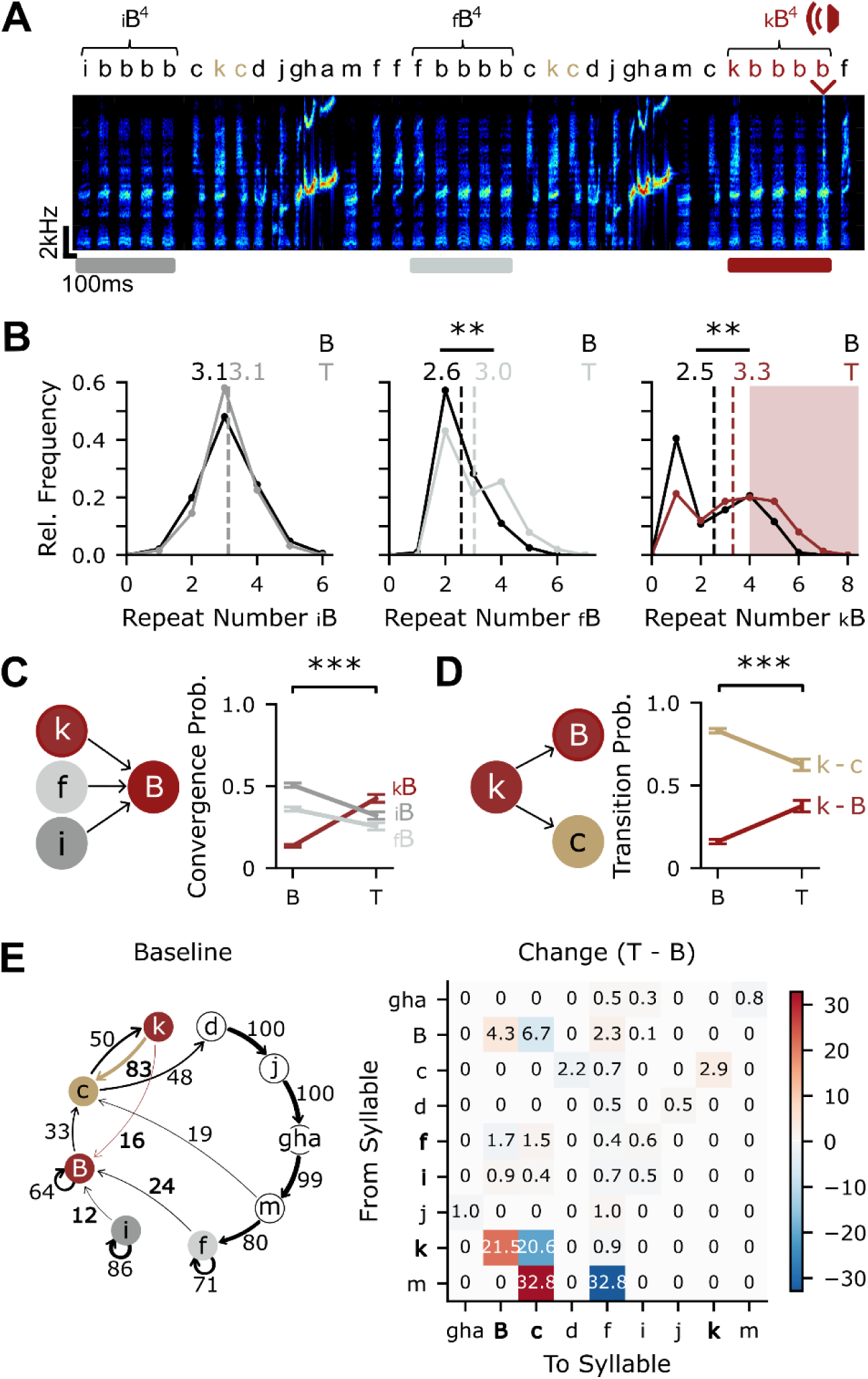
Repeat phrases can be modified in a sequence context-specific way. A) Spectrogram of an example song bout from bird 3. Repeat phrase ‘B’ is marked based on its sequence contexts ‘i’, ‘f’ and ‘k’. The click was played at the fourth and higher positions of sequence context ‘kB’ (red arrowhead). Branch point syllables ‘k-c’ and ‘k-B’ are color-coded. B) Repeat distributions of phrase ‘B’ in the different sequence contexts ‘iB’ (left), ‘fB’ (middle) and ‘kB’ (right) during baseline (B, black) and training (T, color). Dashed lines show mean repeat number. Red shaded region depicts rewarded repeat numbers for the targeted sequence context ‘kB’ (≥ 4). ** = p < 0.01, Mann-Whitney U tests. C) Left: Possible sequence contexts preceding repeat phrase ‘B’. Right: Mean ± s.e.m. convergence probability of ‘B’ for each sequence context in baseline (B) and training (T). *** = p < 0.001, χ^2^-contingency test. D) Left: Branch point at syllable ‘k’. Right: Mean ± s.e.m. transition probability from ‘k’ to each branch in baseline (B) and training (T). *** = p < 0.001, χ^2^-contingency test. E) Left: Transition diagram of song during baseline. Nodes represent syllables and edges show possible transition between syllables. Line thickness and numbers on edges represent transition probabilities in percent. Branch point and context syllables are marked, and respective probabilities are depicted in bold. Probabilities < 5% are not shown. Node ‘gha’ has been condensed for display purposes. Right: Differences in transition probabilities between nodes, showing relative changes between baseline and training (T - B); transition probabilities computed as in the diagram on the left. Red and blue colors depict positive (T > B) and negative (T < B) changes, respectively.

### Learned changes were specific to the repetition of target syllables

The generalization observed in bird 3 made us wonder whether birds might non-specifically increase other repeat phrase types during training, or conversely if increasing the target repeat number might reduce the bird’s ability to maintain long repeat phrases of other phrase types. Figure 4A shows a spectrogram from example bird 4 with multiple repeat or meta-repeat phrases. The increase in repeat number of the targeted meta-repeat ‘HF’ during training was accompanied by increases in repeat phrases ‘E’ and ‘J’, and a significant decrease in repeat number of phrase ‘B’ (Mann-Whitney U tests: all p < 0.01), while ‘D’ did not change (p > 0.05; Fig. 4B). Across birds, we searched for similar generalization across all repeat phrases (n = 30 repeat phrases). While 7/7 target phrases significantly increased their mean repeat number by +24.3% ± 4.5% (see Fig. S2, 2B, G, 3B for all repeat number distributions), only 7/23 non-target phrases significantly increased their mean repeat number, 4/23 non-target phrases significantly decreased their repeat number and the remaining 12 phrases did not exhibit significant changes (mean change in all non-target phrases: +2.2% ± 4.3%). Relative changes in target phrases were significantly larger than those in non-target phrases (Fig. 4C; Mann-Whitney U test target vs. non-target: U = 139.0, p < 0.01).

**Fig. 4:**
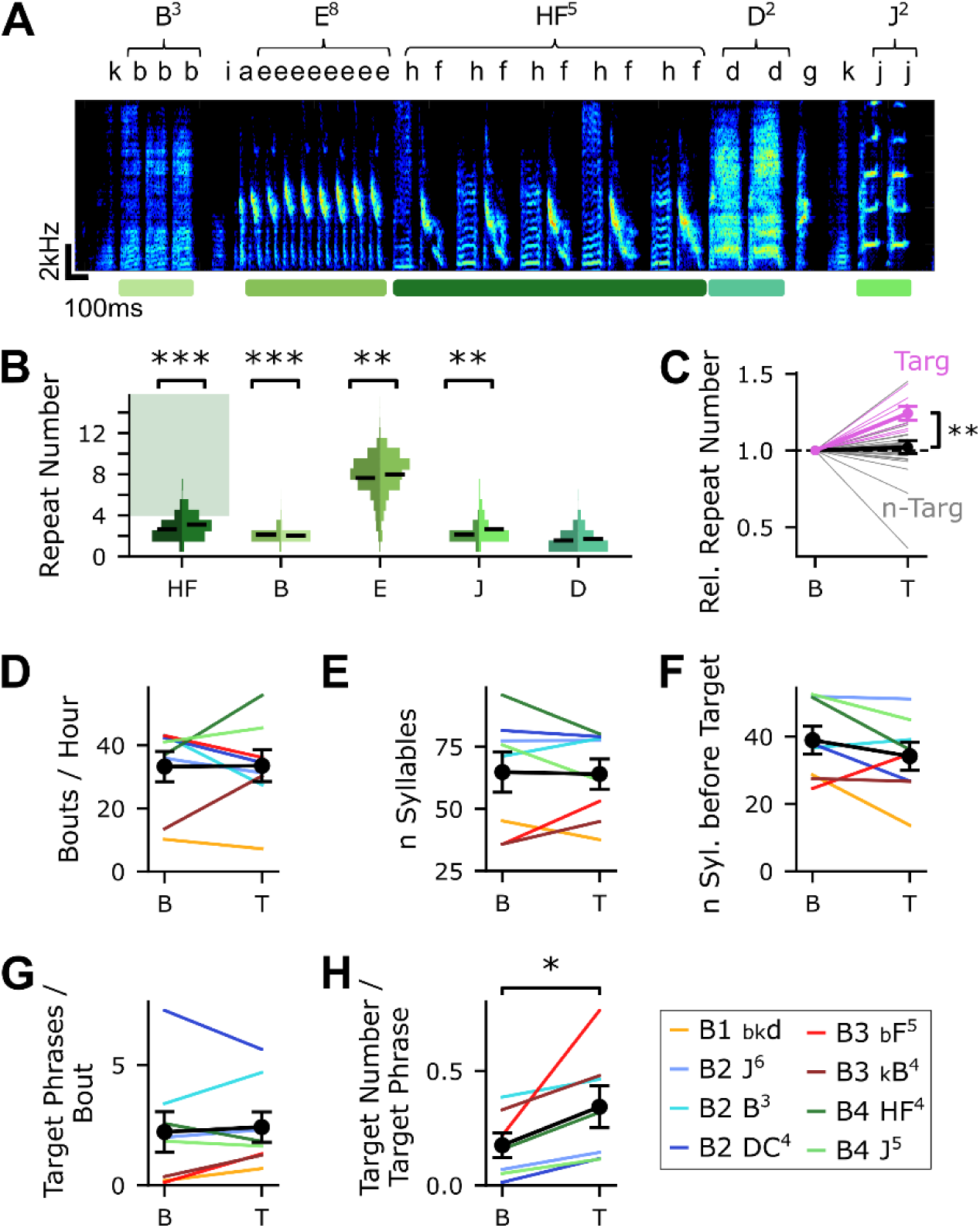
Training-induced changes are specific to targeted repeat numbers. A) Spectrogram of an example song bout from bird 4. Repeat phrases are marked by colored bars, and are depicted by a capital letter, with repeat number given in the exponent. B) Repeat number distributions of each repeat phrase during baseline (left, darker) and training (right, lighter). Extent in the x-direction depicts relative frequency. Black bars indicate mean repeat number. Green shaded area indicates rewarded repeat numbers for the targeted phrase (≥ 4). ** = p < 0.01, Mann-Whitney U tests between baseline and training distributions. C) Relative repeat number of each repeat phrase (pink: targeted repeats, grey: non-target repeats) for all experiments targeting repeat phrases (n = 30 phrases from n = 7 experiments in n = 3 birds). Change during training (T) for each repeat is depicted relative to its normalized baseline value. Thick lines show mean ± s.e.m. change of targeted repeats (pink) and non-targeted repeats (black). ** = p < 0.01, Mann-Whitney U test. D) Mean ± s.e.m. bouts per hour in baseline (B) and training (T, all trials; n = 8 experiments). E) Mean ± s.e.m. number of syllables in baseline (B) and training (T; n = 8 experiments). F) Mean ± s.e.m. number of syllables before the target in baseline (B) and training (T; n = 8 experiments). G) Mean ± s.e.m. number of target phrases per bout in baseline (B) and training (T; n = 8 experiments). H) Mean ± s.e.m. probability of the target phrase containing the target number during baseline (B) and training (T; n = 7 experiments). * = p < 0.05, Wilcoxon signed-rank tests.

To understand whether positive reinforcement led to any changes in motivation to sing or other aspects of singing behavior, we tested whether birds might have made any adjustments to global song structure. We did not find significant differences in song rate (Fig. 4D; baseline 33.3 ± 4.8 bouts/ hour, training 33.5 ± 5.0 bouts/ hour; Wilcoxon signed-rank test: n = 8, z = 17.0, p > 0.05) or mean song length following training (Fig. 4E; baseline 64.8 ± 8.0 syllables, training 64.0 ± 6.1 syllables; Wilcoxon signed-rank test: n = 8, z = 17.0, p > 0.05). If receiving the re-ward on the target syllable was the goal of singing, birds might rearrange their bouts to reach the target syllable earlier. Such a trend towards reducing the number of syllables before the target was observed in some birds, but there was no significant effect at the group level (Fig. 4F; baseline 38.9 ± 4.1 syllables, training 34.2 ± 4.2 syllables; Wilcoxon signed-rank test: n = 8; z = 8.0, p > 0.05). Likewise, there was no difference in the number of times the target phrase appeared within each bout (Fig. 4G; baseline 2.2 ± 0.8 target phrases/ bout, training 2.4 ± 0.6 target phrases/ bout; Wilcoxon signed-rank test: n = 8, z = 13.0, p > 0.05). The only aspect of song that consistently changed across birds and experiments in our analyses was the proportion of target phrases containing the target repeat number or higher (Fig. 4H; baseline 0.2 ± 0.1, training 0.3 ± 0.1; Wilcoxon signed-rank test: n = 7, z = 0.0, p < 0.05). Therefore, birds specifically increased repeat numbers of target repeat phrases, while maintaining all other aspects of song sequencing.

### Song interruptions due to the secondary reinforcer

Learned song changes were analyzed in catch trials, a small percentage of trials in which the secondary reinforcer was omitted. In contrast to changes specific to the target phrase observed in catch trials, most birds showed acute responses to the secondary reinforcer in bouts containing the click sound. Birds frequently interrupted singing once the click was played (Fig. 5A). Figure 5B shows song bouts of example Bird 2 aligned to the target chunk before and after training. This bird decreased the number of syllables following the target significantly (first 20 bouts of baseline 44.3 ± 7.1 syllables, last 20 bouts of training 8.2 ± 5.7 syllables; Mann-Whitney U test: U = 367.5, p < 0.001). Across birds and experiments, there was a significant reduction in the number of syllables following the click playback in training (Fig. 5C; baseline 32.3 ± 5.8 syllables, training 13.8 ± 4.2 syllables; Wilcoxon signed-rank test: n = 8, z = 1.0, p < 0.05). Accordingly, there was a similar reduction in the number of syllables following the target between catch trials, where the bird did not receive feedback or reward, and non-catch trials, where the target was indicated by a click (Fig. 5D; catch trials 29.6 ± 5.3 syllables, non-catch trials 13.8 ± 4.2 syllables; Wilcoxon signed-rank test: n = 8, z = 2.0, p < 0.05). This indicates that most birds prioritized feeding and interrupted their song as soon as they heard the secondary reinforcer. Accordingly, song length (measured in syllables per bout) was lower in non-catch trials than in catch trials (Fig. 5E; mean catch-trails 64.0 ± 6.1, non-catch trials 55.4 ± 7.7; Wilcoxon signed-rank test: n = 8, z = 5.0, p > 0.05).

**Fig. 5:**
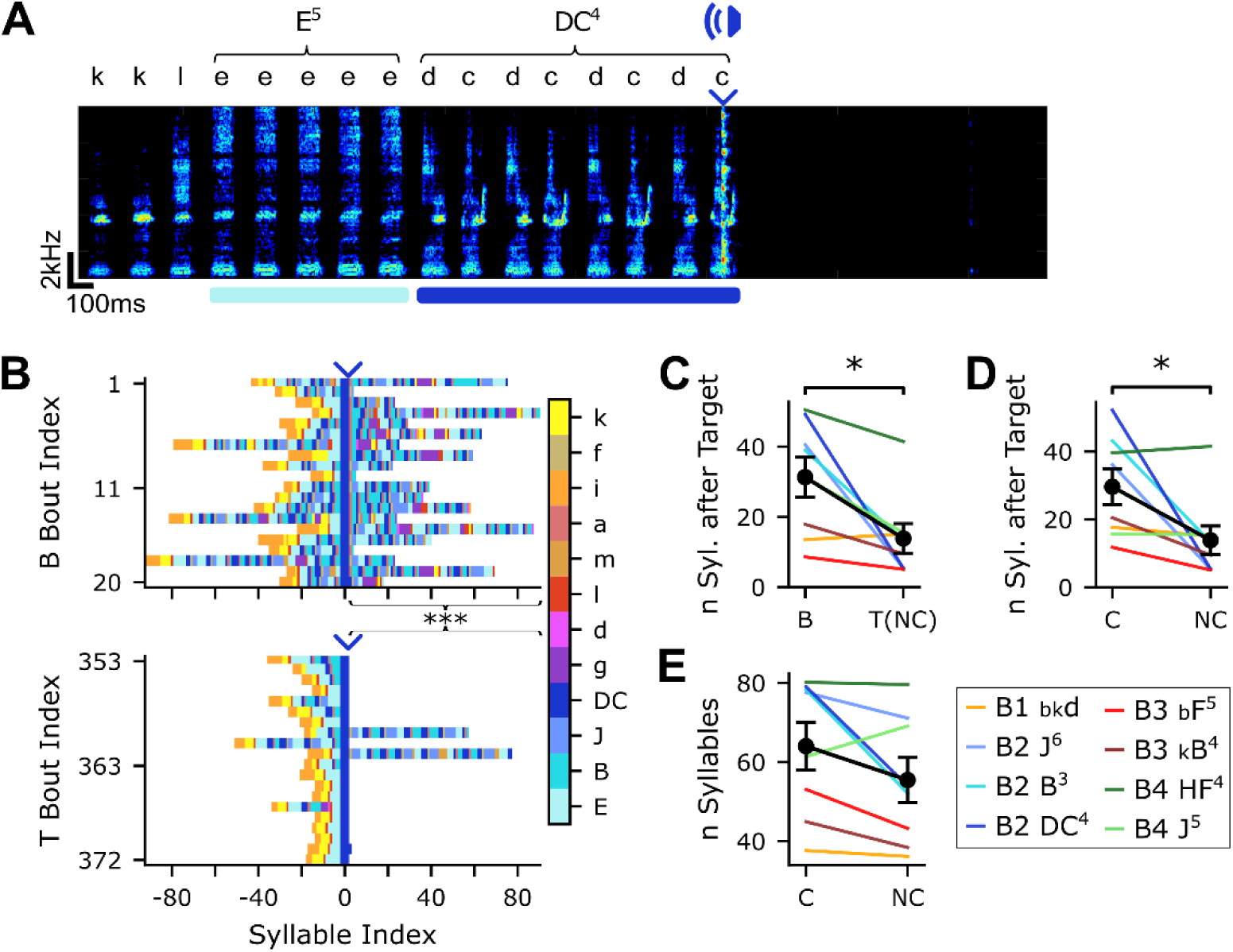
Birds interrupt their song once the click is played. A) Spectrogram of an example song bout from bird 2. A click was played at the fourth position of meta-repeat phrase ‘DC’ (blue arrowhead). B) Syllable sequence of the first 20 bouts of baseline song (top) and last 20 bouts in the training period (bottom) with click playback. Sequences are aligned to the first appearance of the target chunk in each bout (DC^4^; blue arrowhead). Syllables are colored individually, with matching colors for repeat phrases shown in other figures. *** = p < 0.001, Mann-Whitney U test. C) Number of syllables following the target in baseline (B) and non-catch trials (NC, click playback) during training for all experiments (n = 8). Black line shows mean ± s.e.m. * = p < 0.05, Wilcoxon signed-rank test. D) Number of syllables following the target in catch (C) and non-catch (NC) trials for all experiments (n = 8). Black line shows mean ± s.e.m. * = p < 0.05 Wilcoxon signed-rank test E) Number of syllables per bout in catch (C) and non-catch (NC) trials for all experiments (n = 8). Black line shows mean ± s.e.m..

## Discussion

We successfully trained four adult Bengalese finches using a positive reinforcement protocol, pairing a delayed food reward with a click sound as a secondary reinforcer. The click allowed us to target a specific syllable for reward feedback, therefore producing the same temporal precision as experiments using aversive auditory feedback with WN to reduce target transition probabilities [17,34,35]. Different from WN protocols, which do not allow control over which alternative syllable sequence the bird increases, positive reinforcement allows the experimenter-defined targeted increase of specific rare parts of the song. We demonstrate the effectiveness of this protocol for modifying transition probabilities at a branch point, and for modifying repeat number distributions of repeat phrases. These results raise the question of whether the behavioral dynamics for song learning with positive reinforcement resemble the WN protocols which have been studied extensively before, or whether learning to modify song towards food reward follows fundamentally different trajectories. While learning with positive reinforcement can be longer-lasting than with negative reinforcement or punishment [42,43], our learning curves suggest that acquisition as well as return to baseline follow similar dynamics as in WN experiments [17,34,35], though we have not directly compared the effectiveness and stability of both forms of song modification learning in the same birds.

Two other recent studies have shown song modification towards stimuli with positive valence. In deafened zebra finches, visual stimuli that are otherwise perceived as aversive can act as appetitive reinforcers for syllable pitch [44], with dynamics similar to WN feedback for pitch modification learning. One important difference between the study by Zai et al. and our study is that the positive valence of the visual stimulus for the deafened birds was evident in a strong increase in singing rate during training, whereas the birds in our study did not increase singing rate to gain more reward. Instead, they specifically increased repeat numbers to the rewarded level while only minimally adjusting other aspects of their singing behavior (Fig. 4). Bengalese finches receiving pseudo-social reward after the end of song bouts, without direct feedback about the rewarded target, nonspecifically increased repeat numbers to the target in one position of the bout while reducing repeat numbers of the same phrase in other positions in the bout, and showed mixed results for learning at branch points [45]. In contrast, giving the birds direct and specific feedback about the rewarded targets in our study allowed the modification of all occurrences, with comparable effectiveness to WN experiments. Therefore, learning with positive reinforcement can be as effective and specific as WN learning if feedback is given in close temporal contingency to the target. How close the temporal contingency between action and feedback needs to be for effective birdsong modification learning should be investigated in future studies. For pitch modification with WN, delays as short as 100ms have been shown to completely abolish learning [16,20], whereas sequence modification learning seems to be possible with longer delays [34,45].

In some learning tasks, animals exhibit biases or constraints to learn ecologically relevant connections between stimuli or between behavior and rewards. For example, pigeons or hamsters readily learn to display food-related behaviors, but not as easily other behaviors, for food reinforcement [27,46], showing that certain behavior systems are more predisposed to become as-sociated with certain rewards [1,18,28]. Song learning is viewed as an autonomous cognitive module [26,47], meaning an evolutionarily specialised mechanism adapted to solve specific cognitive problems, explaining possible constraints on associations of vocal behavior with food reward [48]. However, song modification learning in response to food rewards described here, as well as to social feedback [45], to visual stimuli in deaf birds [44] and to aversive somatosensory feedback [29] suggest that the song circuit can flexibly incorporate rewarding or punishing feedback from various modalities, and is not inherently specialized for monitoring only auditory feedback [26,29,49]. Likewise, it is not necessary to imitate an ecologically valid social context [45,50] or other behaviorally relevant reinforcer-reward combination [28], but the song system can use an arbitrary learned cue for providing performance-related song feedback. What are the neuronal mechanisms controlling sequence modification through positive reinforcement? Rewarding cues activate dopaminergic midbrain neurons which in all vertebrates project to a wide network of brain areas involved in motivation and reward [4,51–54]. These neurons are involved in reinforcement learning by showing phasic increases or decreases in firing in response to reward-predicting cues and the omission of expected rewards, respectively [55]. In songbirds, a dopaminergic projection to the song-related basal ganglia nucleus Area X encodes ‘performance prediction error’ by responding with phasic increases and decreases in firing to variations in song quality [22,56]. This projection has been shown to be necessary and sufficient to guide learned changes to syllable structure during song modification in adults [23,57–59] and implicated in developmental song learning of syllable structure in juveniles [10,14]. Dopaminergic projections to other areas in the avian brain are involved in other types of learning [54,60–63].

In contrast to learned changes to structure of individual syllables, much less is known about the neuronal mechanism underlying sequence learning in juvenile birds [64–66] or sequence modification learning in adults [17,33–36]. Evidence from lesion and inactivation studies suggests that Area X is involved in controlling syllable repetition [67–70]. In this case, the same dopaminergic midbrain – X connection responsible for syllable structure learning might be involved in reinforcing specific repetition numbers [71]. In the study by Kawaji et al. [45], damage to Area X prevented learning-related changes to syllable repeats, but it is unclear if this represents failure to distinguish between rewarded and non-rewarded trials in their experimental configuration, rather than difficulty controlling repetition itself. Alternatively, transition probabilities between syllables, including self-transitions of repeating syllables, could be stored in the connectivity of premotor networks in song nucleus HVC [72,73,40,74–76]. Specifically, the activity of HVC neurons projecting to Area X is indicative of the total number of repetitions in a repeat phrase [41], so repeat phrases could likely be extended by modifying drive in these HVC neurons [77]. HVC likewise receives dopaminergic input [54,78] that could be involved in reinforcing certain sequences. Finally, non-canonical modulation of Area X by HVC neurons might suggest a close interplay of both areas [14], and potentially other brain areas [79,80] in controlling learned changes to song sequencing.

One important open question is how repeat phrases are represented in the song circuit: Is there only one premotor state associated with the repeating syllable, with one self-transition probability that can be adaptively changed, or could birds learn to increase or decrease self-transitions in a position-specific way? Our results are more consistent with a non-specific increase in self-transition probabilities in all positions before and after the rewarded repeat numbers, suggesting that birds might not learn position-specific changes (Fig. 2E). However, by rewarding all repeat numbers above the target, our experimental design does not allow us to cleanly distinguish between these possibilities. Kawaji et al. [45] omitted reward once repeat phrases became too long, suggesting that it could be possible to learn a specific repeat number, which would not be possible with the simplest form of one-state representation.

Repeat phrases can occur in multiple sequence contexts of preceding syllable types. The sequence context can not only influence repeat distributions at baseline [38,39], but learned modifications can be specific to the rewarded context, with very modest generalization to other contexts (Fig. 3B), similar to partial generalization observed in pitch training [81,82]. Additionally, unrelated repeat phrases of other syllable types could increase or decrease as a result of training (Fig. 4B, C) [38]. This might represent a necessary tradeoff, where the length of one repeat phrase is increased at the cost of decreasing others. Repeating syllables in the wild are performed at or near the performance limit in multiple songbird species [83–85]. However, for undirected song in Bengalese finches, the observed baseline repeat numbers are unlikely to represent a performance limit that would require the bird to make such tradeoffs for increasing repeat numbers beyond the previously occurring distribution. Bengalese finches can increase the number of repeating syllables when singing female-directed song [37], and repetitions can also increase following manipulations or lesions of different song nuclei [67,70,76,77] without specific training [86].

As in previous studies of song modification learning, we analyzed learning performance in catch trials, where feedback is omitted [16,34,35]. Therefore, the observed sequencing changes represent the result of a learning process, and not acute sequence adjustments to the auditory stimulus [32]. While the difference between catch and non-catch trials is typically small for WN experiments, we here found that birds frequently interrupted song immediately after receiving the click. The auditory click itself was likely not perceived as disruptive, because birds never interrupted the syllable receiving the click [87], showed stable song rates and strongly modified transition probabilities towards the click, confirming its appetitive function. There-fore, the birds likely prioritized immediate feeding over singing, once the reward-predicting cue signaled that food would become available [18,88]. It is possible that birds never learned the aspects of our protocol meant to prevent this: the possibility to gain additional reward time by eliciting multiple clicks in one bout, or that the feeder did not become available until after a 1s delay from the end of the bout. Additionally, a strong preference for immediate reward de-livery may have overruled intrinsic motivation to keep singing, representing shifting priorities among competing behaviors [89]. Dopaminergic signals in the basal ganglia respond to cues signaling water reward outside of song, but re-tune to song-related signals during singing [89], representing a context switch in dopamine signaling to the currently prioritized behavior. Our behavioral results suggest that this switch is not absolute [26]: Specifically, birds could both strongly respond to reward cues by interrupting their song in favor of retrieving reward, while at the same time using these cues to modify a specific aspect of song in an adaptive way to increase the probability of receiving reward cues in future renditions. Interestingly, the reward prediction of the auditory cue seems to also overrule monitoring of auditory feedback to assess song performance, resembling the positive valence of female calls which are likewise ‘disturbing’ auditory feedback of song [89,90].

In laboratory settings, some animals can learn to instrumentalize vocalization, producing or withholding calls for food or water rewards [91–97], including the production of a specific number of calls [98]. We have shown that specific features within learned birdsong similarly can be shaped to obtain food reward. This allows the experimentally targeted increase of specific song features in adult birds, and demonstrates that plasticity in the song system has access to general cognitive processes and can to some extent be instrumentalized in the pursuit of external goals [34,99].

## Material and Methods

### Animals

Five adult (age 471 - 651 days) male Bengalese finches (*Lonchura striata domestica*) were used in the experiments, each participating in one to three training sessions. Birds that participated in multiple trainings were returned to group housing in aviaries in between experiments. All birds were bred and raised in the lab. For the duration of the experiment, birds were housed in cages inside sound-attenuating chambers (120×50×50cm) under a 14:10 hour light cycle at 25°C and 50% humidity with *ad libitum* food, water and standard enrichment (sand baths, nests, fresh greens). During the conditioning and song-training phases, the feeder was activated for several hours a day, and birds were restricted to a part of the cage with food provided only from the automated feeder. Outside of feeder activation hours in the mornings and evenings, birds were housed with food *ad libitum* and fresh greens and accompanied by a social partner. Birds participating in the experiments were weighed regularly to ensure that weight remained stable and monitored with a webcam (BRIO 4K PRO, Logitech). All experiments were performed in accordance with animal protocols approved by the Regierungspräsidium Tübingen, Germany.

### Experimental setup

Songs were recorded using the custom software Moove [36] with either a Rode M5 MP micro-phone (Australia) or an omnidirectional condenser microphone (AT803, audiotechnica) connected to a Steinberg UR12/IXO12 audio interface (Germany) and a recording computer. An Arduino (Uno R3) with an attached relay shield (ARD SHD RELAY V3 Arduino shield - Re-lais V3) was connected to a speaker (BBCZ speaker for Micro:Bit; WAVESHARE) and a motor (Linear motion actuator, 30mm/s 20N DC 12V, Hilland) controlling a custom-made feeder (similar to [100]). The feeder could be moved by 3cm into and out of the experimental cage, where the feeder tray containing a standard seed mix was positioned centrally between a u-shaped perch. A tone signal (3ms duration, 500Hz), created by the built-in arduino function tone, was utilized to play the click sound with the speaker.

### Conditioning procedure

All birds participated in a conditioning procedure prior to song training, where the click sound was paired with feeder movement into the cage (delay of 800ms). The frequency and duration of feeder activation was successively reduced over multiple conditioning days, depending on each bird’s reliability and latency to approach the feeder. In the final stage, the feeder was activated sparsely and pseudo-randomly over the day, and birds had learned to approach the feeder immediately after the click sound.

### Song training

The song training for each experiment included two days of baseline recordings, six training days and two post-baseline recordings, one immediately following training and another after approximately one week. The song training phase lasted approximately eight hours per day. During this phase, the click sound was played when the target syllable was detected. Between 5 - 10% of songs were designated as catch trials with sound playback and reward omitted, in order to distinguish learning-related effects, which also occur on catch trials, from acute responses to the sound playback. To measure targeting accuracy, we calculated correct hits, false positives and misses of playback after manual offline correction. In these experiments, Moove online-classification achieved a mean hit rate of 84.6 ± 4.5% (Fig. S1, n = 9), with most false positive targets occurring on the target syllable in other repeat positions. For bird 5, targeting accuracy was below 50% for several training days, thus we excluded this bird from analyses because the conditions for successful learning were not provided. For each bird, the target syllable was chosen as a rare branch following a branch point (n = 1) or rare repeat numbers that occurred in baseline screening with a mean frequency of 11.5 ± 4.5% (range: 1.4% to 38.5%, n = 8). Whenever a hit was detected, the feeder moved inside the cage after 1s silence from the end of the bout. The duration of feeder presentation was contingent upon the number of clicks accumulated in the bout. The minimum duration for one click was set individually for each training, based on estimated target occurrence to ensure sufficient food access (mean 23.1s ± 3.4s, range 10 - 42s, n = 9) and each additional click increased this duration (mean +9.2 ± 0.6s, range +5 - +10s, n = 9).

### Song analysis

As in other studies, we defined a song bout as a continuous period of singing separated by 2s of silence [33,101]. We used Moove [36] to train neural networks on baseline recordings for every bird, enabling online classification during song. To check the accuracy for classification of the target sequence, a subset of bouts was manually checked on each training day. Annotation of classified syllables was semi-automatically corrected offline before performing analysis on the data. The syllable sequence was then analyzed as described in detail in Koparkar et al. [33]. One syllable branching into two or more distinct syllables in a probabilistic manner was defined as a branch point [32,33]. The same syllable occurring repeatedly in sequence was defined as a repeat phrase [31,33,37,38] and is depicted as a capital letter in spectrogram examples and transition diagrams. Chunks were defined as a syllable sequence that occurs in a fixed order [102]. We defined a meta-repeat as a repeating chunk, i.e. a repeat phrase of variable duration repeating a short sequence of syllables in fixed order [103]. Song sequencing at the end of training was quantified during the last three days of training (T4 - 6). Training values (T) were analyzed based on catch-trials (C) only, unless stated otherwise.

### Statistical analysis

All analyses were performed using custom-written python scripts (version 3.12) and open-source statistic packages. All analyses include eight experiments (n = 8) from four birds unless stated otherwise. All analyses of repeat number include 7 experiments from three birds (birds 2 - 4) which received targeting at repeat phrases.

Mean repeat numbers and transition probabilities were calculated by pooling all occurrences, without averaging over song bouts or days. Mean values are given including the standard error of the mean (s.e.m.). We compared the group mean of baseline and training or catch and non-catch conditions using Wilcoxon signed-rank tests with α = 0.05, treating each training as an individual data point. To assess differences in branchpoint probabilities between baseline and training or within meta-repeats, we conducted χ^2^-contingency tests with α = 0.05. If one cell in the contingency table had an expected frequency < 5, we conducted a Fisher’s exact test instead. Differences between repeat distributions in baseline and training were tested using a Mann-Whitney U test, α = 0.05, preceded by Shapiro-Wilk tests revealing almost exclusively non-parametric distributions. Changes in self-transition probabilities at different positions within repeats were assessed by computing mean self-transition probability over all trainings, creating one baseline and training value for each position. A Wilcoxon signed-rank test was performed on mean values for each position. Differences in training-induced change between target and non-target repeats were assessed by comparing the distributions of relative change using a Mann-Whitney U test. All p-values were Holm-corrected for multiple comparisons.

## Acknowledgements

This work was supported by the Deutsche Forschungsgemeinschaft (DFG – Project number 536953998) to L.V. and a PhD fellowship by the Ersatzausschreibung Landesgraduiertenförderung by the University of Tübingen to F.H. We thank members of the Veit lab Avani Koparkar, Jacqueline Göbl, Lioba Fortkord, Priya Binwal and Abhilipsa Das for helpful discussions and comments on the manuscript, and Nils Riekers for advice on the setup.

## Supplementary Figures

**Fig. S1:**
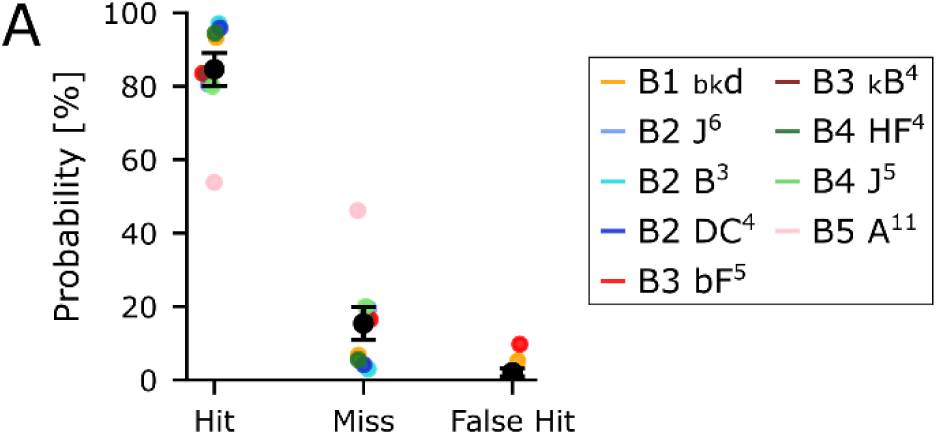
Targeting accuracy for each experiment. Probability [%] of each target to be a hit (correct playback) or miss (missing playback), and probability of false hits (playback at non-target positions of the phrase) for all experiments (n = 9) in all birds (n = 5). Black bar shows mean ± s.e.m. across experiments.

**Fig. S2:**
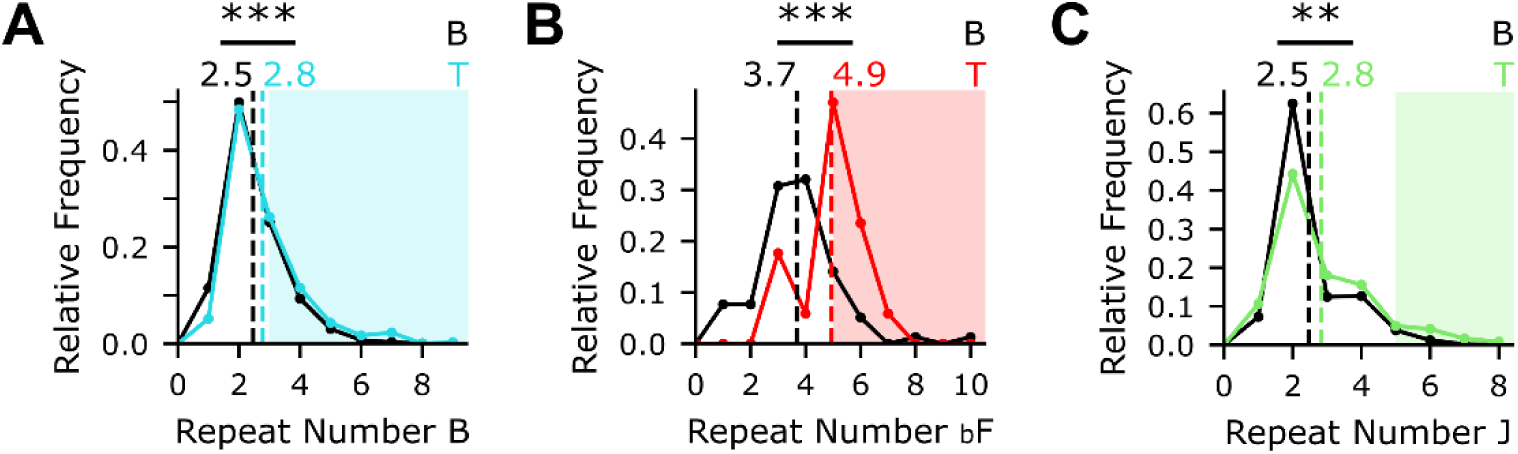
Repeat distributions for remaining experiments. Repeat distributions for baseline (black) and training (color) of repeat phrase ‘B’ in bird 2 (A), repeat phrase ‘F’ in sequence context ‘bF’ in bird 3 (B) and repeat phrase ‘J’ in bird 4 (C). Dashed lines show mean repeat number. Shaded regions depict rewarded repeat numbers: ≥ 3 (A), ≥ 5 (B) and ≥ 5 (C), respectively. ** = p < 0.01, *** = p < 0.001, Mann-Whitney U tests.

